# Quality control and processing of nascent RNA profiling data

**DOI:** 10.1101/2020.02.27.956110

**Authors:** Jason P. Smith, Arun B. Dutta, Kizhakke Mattada Sathyan, Michael J. Guertin, Nathan C. Sheffield

**Affiliations:** Center for Public Health Genomics, University of Virginia; Department of Public Health Sciences, University of Virginia; Department of Biomedical Engineering, University of Virginia; Department of Biochemistry and Molecular Genetics, University of Virginia

## Abstract

Nascent RNA profiling is growing in popularity; however, there is no standard analysis pipeline to uniformly process the data and assess quality. Here, we introduce PEPPRO, a comprehensive, scalable workflow for GRO-seq, PRO-seq, and ChRO-seq data. PEPPRO produces uniformly processed output files for downstream analysis and assesses adapter abundance, RNA integrity, library complexity, nascent RNA purity, and run-on efficiency. PEPPRO is restartable and fault-tolerant, records copious logs, and provides a web-based project report. PEPPRO can be run locally or using cluster, providing a portable first step for genomic nascent RNA analysis.

**Availability:** BSD2-licensed code and documentation: https://peppro.databio.org.

## Background

Steady-state transcription levels are commonly measured by RNA-seq, but there are many advantages to quantifying *nascent* RNA transcripts: First, it measures the transcription process directly, whereas steady-state mRNA levels reflect the balance of mRNA accumulation and turnover. Second, nascent RNA profiling measures not only RNA polymerase occupancy, but also orientation by default, whereas traditional RNA-seq requires specific library preparation steps to capture orientation. Third, nascent RNA profiling measures unstable transcripts, which can be used to infer regulatory element activity and identify promoters and enhancers *de novo* by detecting bidirectional transcription and clustered transcription start sites (TSSs)^1,2^. Fourth, nascent RNA profiling can be used to determine pausing and RNA polymerase accumulation within any genomic feature. These advantages have led to growing adoption of global run-on (GRO-seq), precision run-on (PRO-seq), and, most recently, chromatin run-on (ChRO-seq) experiments^3–5^. With increasing data production, we require analysis pipelines for these data types. While tools are available for downstream analysis, such as to identify novel transcriptional units and bidirectionally transcribed regulatory elements^1,6–10^, there is no comprehensive, unified approach to initial sample processing and quality control.

Here, we introduce PEPPRO, an analysis pipeline for uniform initial sample processing and novel quality control metrics. PEPPRO features include: 1) a serial alignment approach to remove ribosomal DNA reads; 2) nascent transcription-specific quality control outputs; and 3) a modular setup that is easily customizable, allowing modification of individual command settings or even swapping software components by editing human-readable configuration files. PEPPRO is compatible with the Portable Encapsulated Projects (PEP) format, which defines a common project metadata description, facilitating interoperability^11^. PEPPRO can be easily deployed across multiple samples either locally or via any cluster resource manager, and we also produced a computing environment with all the command-line tools required to run PEPPRO using either docker or singularity with the bulker multi-container environment manager^12^. Thus, PEPPRO provides a unified, cross-platform pipeline for nascent RNA profiling projects.

## Results

### Pipeline overview and data description

PEPPRO starts from raw, unaligned reads, and produces a variety of output formats, plots, and quality control metrics. Briefly, pre-alignment steps include removing adapters, deduplicating, trimming, and reverse complementation (Fig. 1). PEPPRO then uses a serial alignment strategy to siphon off unwanted reads from rDNA, mtDNA, and any other user-provided decoy sequences. It aligns reads and produces signal intensity tracks as both single-nucleotide counts files and smoothed normalized profiles for visualization. PEPPRO also provides a variety of plots and statistics to assess several aspects of library quality, such as complexity, adapter abundance, RNA integrity and purity, and run-on efficiency (See Methods for complete details).

**Fig. 1:**
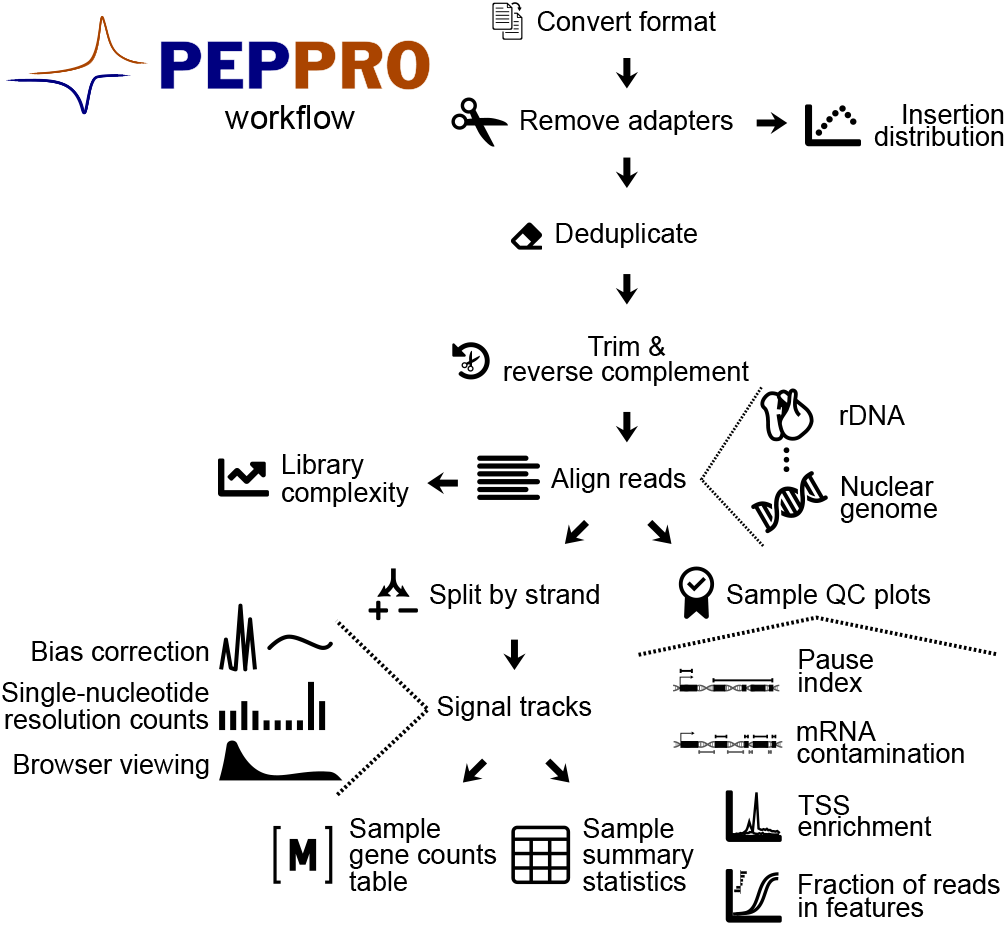
PEPPRO steps for genomic run-on data. PEPPRO starts from raw sequencing reads and produces a variety of quality control plots and processed output files for more detailed downstream analysis.

To evaluate PEPPRO on different library types, we assembled a test set of run-on libraries with diverse characteristics (Fig. 2A). Our test set includes 7 previously published libraries: 2 ChRO-seq, 2 GRO-seq, and 3 PRO-seq^5,13–15^. We ran each of these samples through PEPPRO as a test case and visualized the data in a genome browser (Fig. 2B). To demonstrate PEPPRO’s setup for differential expression analysis, we also generated paired-end PRO-seq libraries from H9 cell culture samples either naive or treated with romidepsin, a histone deacetylase inhibitor (HDACi). This test set therefore provides a range of qualities, protocols, and issues, providing a good test case for demonstrating the novel quality control features of PEPPRO and how to distinguish high-quality samples.

**Fig. 2:**
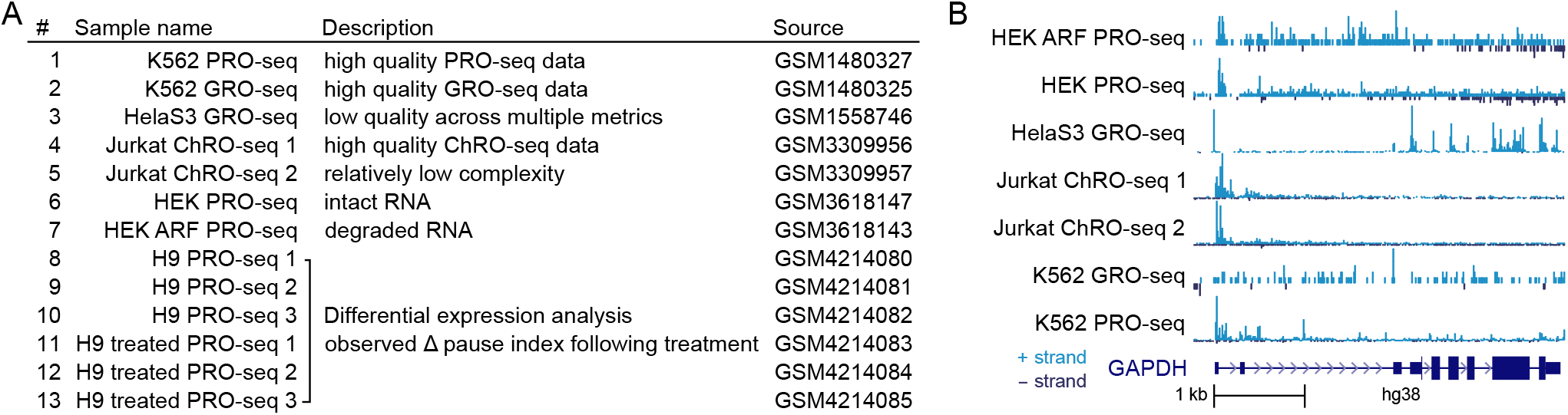
PEPPRO test set data table and signal tracks. A) Table showing the attributes of samples collected for our test set. Complete metadata is available from the PEPPRO website. B) Read count normalized signal tracks from published data are visualized within a browser.

To demonstrate how PEPPRO responds to mRNA contamination, we also generated a set of 11 samples built from a single PRO-seq library (GSM1480327) that we spiked with increasing amounts of RNA-seq data(GSM765405) (Fig. S1). We ran PEPPRO on our public test set, our differential expression test set, and our spike-in set. Results of PEPPRO can be explored in the PEPPRO HTML-based web report, which displays all of the output statistics and QC plots (see PEPPRO documentation). Here, we describe each plot and statistic produced by PEPPRO.

### Adapter ratio

A common source of unwanted reads in PRO/GRO/ChRO-seq libraries results from adapter-adapter ligation. These methods require two independent ligation steps to fuse distinct RNA adapters to each end of the nascent RNA molecule. The second ligation can lead to adapter-adapter ligation products that are amplified by PCR. The frequency of adapter-adapter ligation can be reduced by molecular techniques (see Methods), but these are not always possible and many experiments retain adapters in high molar excess, leading to substantial adapter-adapter sequences.

PEPPRO counts and reports the fraction of reads that contain adapter-adapter ligation products, then removes adapter sequences and adapter-adapter ligation sequences before downstream alignment. In our test, all samples had fewer than 50% adapter-adapter ligation reads (Fig. S2). Higher rates do not necessarily reflect lower quality samples, but rather indicate a suboptimal ratio of adapters during the library preparation or exclusion of the gel extraction size selection step. Excess adapters indicate that future sequencing will be less informative, leading to increased depth requirements, and therefore inform on whether to sequence a library deeper, tweak the adapter ratio in future samples, or include a size selection step. In our hands, we aim for adapter-adapter ligation abundance between 20-50% with no size selection step, or less than 5% if the final library is polyacrylamide gel electrophoresis (PAGE) purified. Libraries with no adapter-adapter ligation indicate that size selection was too stringent, and may actively select against short RNA insertions from specific classes of nascent RNA, such as RNAs from promoter-proximal paused polymerases^16^.

### RNA integrity

A common indicator of RNA sample quality is the level of RNA integrity. RNA integrity can be assessed by plotting the distribution of RNA insert sizes, which will be smaller when RNA is degraded. For a highly degraded library, we expect insert sizes below 20 nucleotides, which corresponds to the length of RNA between the RNA polymerase exit channel and 3^*I*^ RNA end. These nucleotides are sterically protected from degradation^17^, so high frequency of insert sizes below 20 indicates that degradation occurred after the run-on step^5^ (Fig. 3A).

**Fig. 3:**
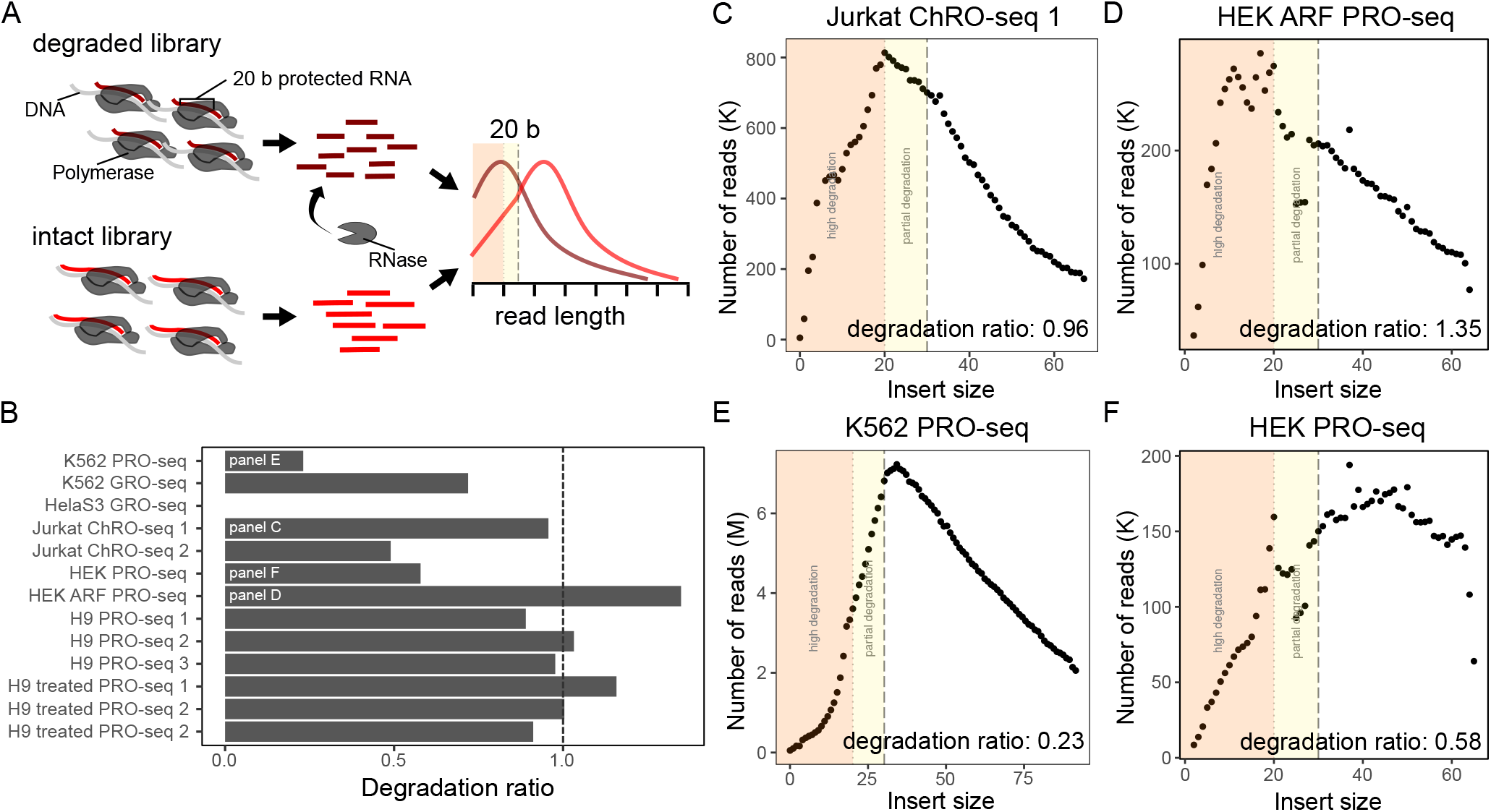
RNA integrity is assessed with degradation ratios and insert sizes. A) Schematic illustrating intact versus degraded libraries. B) Degradation ratio for test samples (HelaS3 GRO sample could not be calculated; Values less than dashed line (1.0) are considered high quality). C-F) Insert size distributions for: C, a degraded single-end library; D, a degraded paired-end library; E, a non-degraded single-end library; and F, a non-degraded paired-end library (orange shading represents highly degraded reads; yellow shading represents partially degraded reads).

PEPPRO uses a novel method to calculate the insert size distribution that applies to both single- and paired-end data (see Methods). PEPPRO reports the ratio of insert sizes from 10-20 nucleotides versus 30-40 nucleotides, which measures RNA integrity because more degraded libraries have higher frequency of reads of length 10-20, whereas less degraded libraries have more reads of length 30-40. Using our test set, we found that PRO-seq libraries with a ratio *<* 1 should be considered high quality (Fig. 3B). A single-end ChRO-seq library that was intentionally degraded with RNase prior to the run on step^5^ has a degradation ratio near 1 with a insertion distribution plot showing a peak at 20 nucleotides (Fig. 3C). A poor quality paired-end PRO-seq library contains many RNA species falling within the 10-20 range (Fig. 3D). High-quality libraries show plots that peak outside of the sub-20-nucleotide degradation zone (Fig. 3E, F).

### Library complexity

Library complexity measures the uniqueness of molecules in a sequencing library (Fig. 4A). For conventional RNA-seq, shearing is random, so paired-end reads with the same start and end coordinates may be assumed to be PCR duplicates. In contrast, in PRO-seq, transcription start sites account for many of the 5′ RNA ends, and promoter proximal pause sites can focus the 3′ end of the RNA^4^, so independent insertions with the same end points are not necessarily PCR duplicates. As a result, unfortunately, this means it is not possible to calculate complexity generally.

**Fig. 4:**
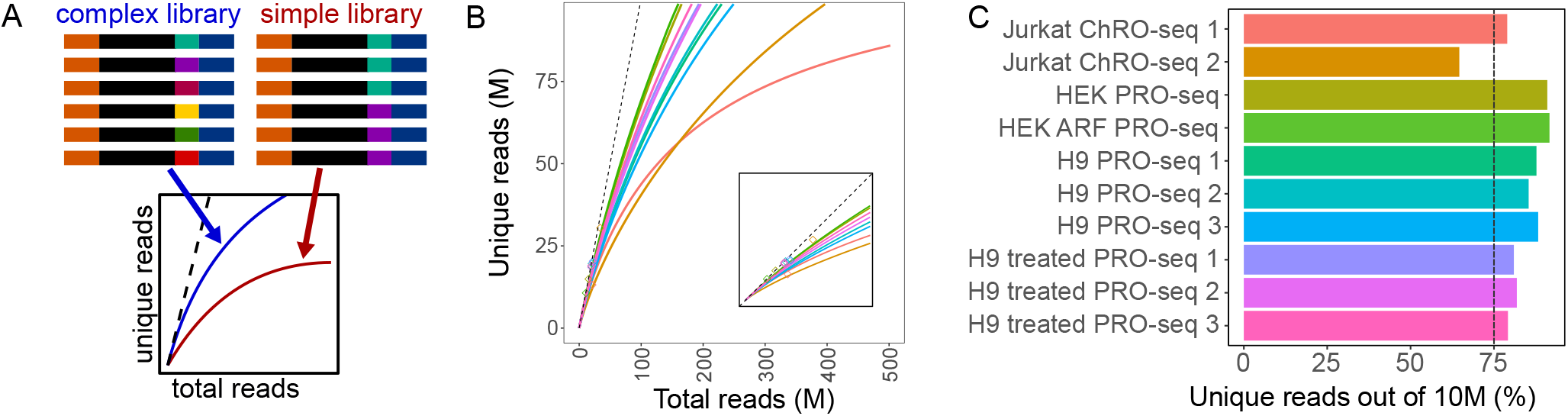
Library complexity is measured with unique read frequency distributions and projections. A) Schematic demonstrating PCR duplication and library complexity (dashed line represents completely unique library). B) Library complexity traces plot the read count versus externally calculated deduplicated read counts. Deduplication is a prerequisite, so these plots may only be produced for samples with UMIs. Inset zooms to region from 0 to double the maximum number of unique reads. C) The position of curves in panel B at a sequencing depth of 10 million reads (dashed line represents minimum recommended percentage of unique reads).

Recent PRO-seq protocols resolve this by incorporating a unique molecular identifier (UMI) into the 3′ adapter, which PEPPRO uses to distinguish between PCR duplicates and independent RNA molecules with identical ends. For data that includes UMIs, PEPPRO accommodates multiple software packages for read deduplication, including `seqkit`^18^ and `fqdedup`^19^. PEPPRO calculates library complexity at the current depth, reporting the percentage of PCR duplicates. In our test samples, we found that libraries with at least 75% of reads unique at a sequencing depth of 10 million can be considered high quality (Fig. 4C). PEPPRO also invokes preseq^20^ to project the unique fraction of the library if sequenced at higher depth (Fig. 4B). These metrics provide a direct measure of library complexity and allow the user to determine value of additional sequencing. However, because nascent RNA reads cannot be effectively deduplicated using the standard approach applied to traditional RNA-seq, complexity metrics are only calculated for samples with UMIs.

### Nascent RNA purity

One challenge specific to nascent RNA sequencing is ensuring that the library targets nascent RNA specifically, which requires eliminating the more abundant processed rRNA, tRNA, and mRNA transcripts. Early run-on protocols included 3 successive affinity purifications, resulting in 10,000-fold enrichment over mRNA and over 98% purity of nascent RNA^3,4^. Newer run-on protocols recommend fewer affinity purifications^15^. Therefore, assessing the efficiency of nascent enrichment is a useful quality control output.

To estimate the *nascent purity* of RNA, PEPPRO provides two results: an mRNA contamination metric and a rDNA alignment rate. First, PEPPRO assesses nascent RNA purity by calculating the exon to intron read density ratio (Fig. 5A). A nascent RNA sequencing library without polymerase pausing would have a ratio of exon density to intron density of ≈ 1. Because promoter-proximal pausing inflates this ratio, PEPPRO excludes the first exon from this calculation. In our test samples, the median exon-intron ratio is between 1.0 and 1.8 for high quality libraries (Fig. 5B). Our *in silico* spike-in of conventional RNA-seq increases this ratio proportionally to the level of mRNA contamination (Fig. 5B). This ratio varies substantially among genes and PEPPRO produces histograms to compare in more detail among samples (Fig. 5C-F). By comparing these values to the spike-in experiment, we can estimate the level of mRNA contamination of a library (Fig. 5E, F). A second measure of nascent purity is to evaluate relative rRNA abundance.

**Fig. 5:**
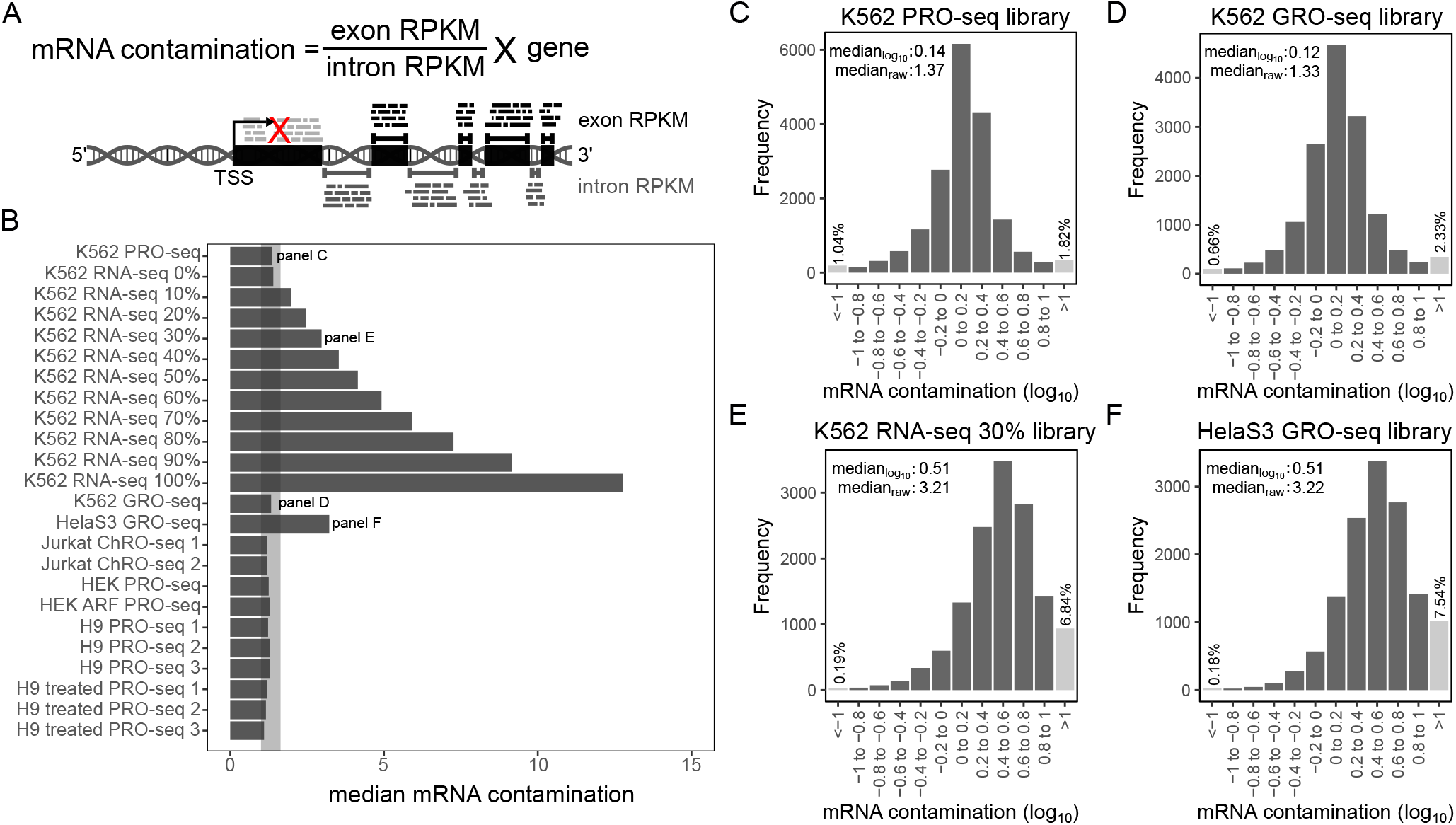
Nascent RNA purity is assessed with the exon-intron ratio. A) Schematic demonstrating mRNA contamination calculation. X represents the exclusion of the first exon in the calculation. B) Median mRNA contamination metric for test set samples (Shaded region represents recommended range (1-1.8)). C) Histogram showing the distribution of mRNA contamination score across genes in the K562 PRO-seq sample. D) As in panel C for a GRO-seq library. E) mRNA contamination distribution for K562 PRO-seq spiked with 30% K562 RNA-seq. F) mRNA contamination distribution for HelaS3 GRO-seq is comparable to the 30% RNA-seq spike-in sample.

Since rRNA represents the vast majority of stable RNA species in a cell, overrepresentation of rRNA reads indicates poor nascent RNA enrichment. We find that high-quality nascent RNA libraries typically have less than 20% rRNA alignment (Fig. S3). In contrast, between 70% and 80% of mature RNA in a cell is rRNA. Therefore, the ratio of rRNA aligned reads compared to the all other reads reflects mature RNA contamination. To demonstrate, we calculated the correlation between the exon-intron read density ratio and the rRNA-to-aligned-reads ratio using the primary set of test samples with additional samples (GSE126919) to increase power. We found these two measures are significantly correlated (Fig. S4). Exon-intron read density ratio is a more robust measure of nascent RNA purity, as the fraction of nascent rRNA transcription is likely to be distinct among cell lines. However, PEPPRO still reports the rDNA alignment ratio as an orthogonal measure of nascent RNA purity and overall library quality.

### Run-on efficiency

Another quality metric for run-on experiments is run-on efficiency. Typically, gene-body polymerases extend efficiently during the nuclear run-on step, but promoter-proximal paused polymerases require either high salt or detergent to do so^21,22^. Because these treatments vary, leading to varying run-on efficiency, PEPPRO employs two methods to assess run-on efficiency: *pause index* and *TSS enrichment*. First, we define the pause index as the ratio of the density of reads in the *pausing* region versus the density in the corresponding gene body (Fig. 6A; see Methods). PEPPRO plots the frequency distribution of the pause index across genes. A greater pause index indicates a more efficient run-on, as a higher value indicates that paused polymerases efficiently incorporate the modified NTPs. As test of this metric, we analyzed GRO-seq data that was generated in the presence and absence of the anionic detergent Sarkysol^22^. Paused polymerases necessitate detergent to run on and incorporate NTPs efficiently, thus the pause index drops substantially in the absence of Sarkysol (Fig. 6B,C). We found in our test samples that an efficient run-on process has a median pause index greater than 10 (Fig. 6D). For more detail, PEPPRO produces frequency distribution plots that show an exponential distribution among genes for an efficient library (or a normal distribution on a log scale, Fig. 6E) and a shifted distribution for an inefficient run-on (Fig. 6F).

**Fig. 6:**
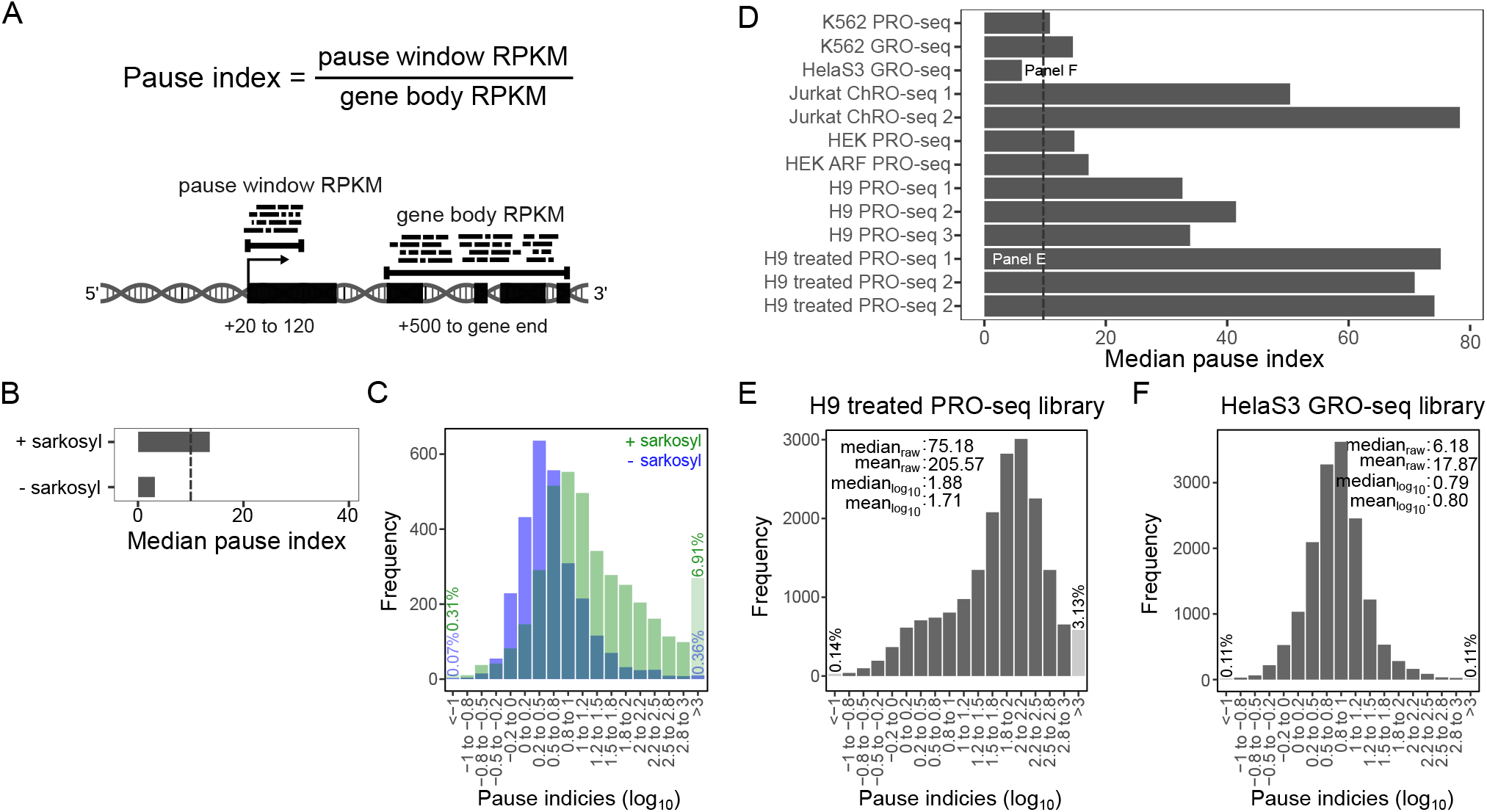
Run-on efficiency is measured with pause indices. A) Schematic demonstrating pause index calculation. B) Pause index values for Drosophila melanogaster GRO-seq libraries with (GSM577247) or without sarkosyl (GSM577248). C) The histogram of pause index values is shifted to the right upon addition of sarkosyl in GRO-seq libraries. D) Pause index values for test set samples (Values above the dashed line are recommended). E) High pause index identified in H9 treated PRO-seq. F) Low pause index from HelaS3 GRO-seq.

As a second assessment of run-on efficiency, PEPPRO aggregates sequencing reads at TSSs to plot and calculate a TSS enrichment score. PEPPRO plots aggregated reads 2000 bases upstream and downstream of a reference set of TSSs. The normalized TSS enrichment score is calculated by taking the average base coverage in a 100 bp window around the peak divided by the average coverage in the first 200 bases. Efficient TSS plots show a characteristic PRO-seq pattern with an upstream peak for divergently transcribing polymerases and a prominent peak representing canonical paused polymerases (Fig. S5). PEPPRO also summarizes these values across samples.

### Read feature distributions

PEPPRO also produces plots to visualize the *fraction of reads in features*, or FRiF. The cumulative FRiF (cFRiF) plot provides an information-dense look into the genomic distribution of reads relative to genomic features. This analysis is a generalization of the more common *fraction of reads in peaks* (FRiP) plots produced for other data types^23^ with two key differences: First, it shows how the reads are distributed among different features, not just peaks; and second, it uses a cumulative distribution to visualize how quickly the final read count is accumulated in features of a given type. To calculate the FRiF, PEPPRO overlaps each read with a feature set of genomic annotations, including: enhancers, promoters, promoter flanking regions, 5′ UTR, 3′ UTR, exons, and introns (Fig. 7). The individual feature elements are then sorted by read count, and for each feature, we traverse the sorted list and calculate the cumulative sum of reads found in that feature divided by the total number of aligned reads. We plot the read fraction against the *log*_10_ transformed cumulative size of all loci for each feature. This allows the identification of features that are enriched for reads with fewer total features and total genomic space. Additionally, PEPPRO calculates the non-cumulative FRiF by taking the *log*_10_ of the number of observed bases covered in each feature over the number of expected bases in each feature to identify enriched genomic features (Fig. 7).

**Fig. 7:**
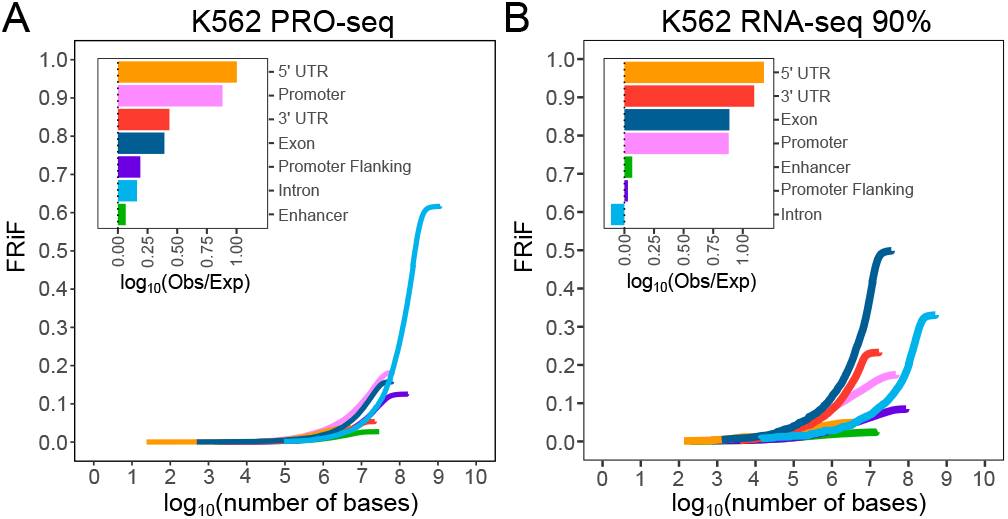
Fraction of reads in genomic features. A) K562 PRO-seq represents a “good” cumulative fraction of reads in features (cFRiF) and fraction of reads in features (FRiF) plot. B) K562 PRO-seq with 90% K562 RNA-seq spike-in represents a “bad” FRiF/PRiF.

In our test samples, high-quality libraries have a characteristic pattern with slow accumulation but high total of reads in introns, and fast accumulation but lower total of reads in promoter elements. ChRO-seq libraries have an increased promoter emphasis and higher mRNA contamination indicated by an increase in reads in promoters and exons at the cost of reads in introns and promoter flanking regions (Fig. S6). Additionally, the RNA-seq spike-in samples demonstrate the increasing prevalence of exonic reads and 3′ UTR at the cost of intronic sequences (Fig. S7). These plots are therefore a useful general-purpose quality control tool that reveal substantial information about a sample in a concise visualization.

### Differential expression

The focus of PEPPRO is in the pre-processing relevant for any type of biological project. The output of PEPPRO sets the stage for downstream analysis specific to a particular biological question. Perhaps the most common downstream application of nascent transcription data is differential expression analysis. PEPPRO allows the user to easily run a differential comparison using dedicated software like the DESeq2 bioconductor package^24^. To demonstrate this, we included PRO-seq libraries from H9 human cutaneous T-cell lymphoma cell lines treated with either DMSO (n=3) or an HDAC inhibitor (n=3).

To facilitate differential expression analysis, PEPPRO produces a project-level counts table that may be loaded in R using `pepr`, and, in a few lines of code, converted quickly into DEseq data sets ready for downstream DESeq analyses (See Supplemental text). Using this approach, we ran a differential expression analysis comparing romidepsin-treated against untreated samples (Fig. 8A). We identified many genes with significantly different read coverage. As an example, the PTPN7 gene showed clear differences in counts (Fig. 8B), which we can further visualize using the browser track outputs generated by PEPPRO (Fig. 8C). This analysis demonstrates how simple it is to ask a downstream biological question starting from the output produced by PEPPRO.

**Fig. 8:**
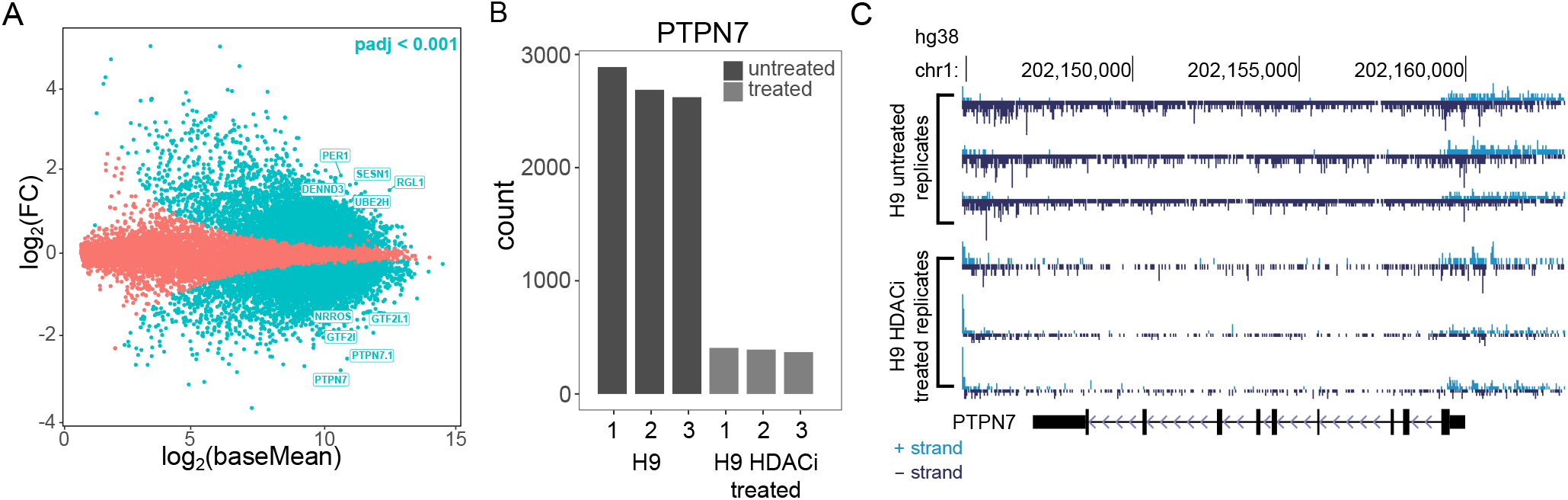
Differential analysis with the PEPPRO counts matrix. A) MA plot between H9 DMSO versus H9 200nM romidepsin treated PRO-seq libaries (dots = genes; top 10 most significant genes labeled; n=3/treatment). B) Most significantly differential gene count differences. C) Read count normalized signal tracks from the differential analysis.

### Metric robustness

To evaluate the robustness of our metrics across sequencing depth and library complexity, we ran PEPPRO on subsampled single-end and paired-end with UMI PRO-seq libraries (Fig. S8, Fig. S9). Our metrics remained constant across sequencing depth from as few as 10M reads to well over 100M (Fig. S10, Fig. S11). We also generated synthetic low complexity paired-end with UMI PRO-seq libraries and our metrics remain robust to reductions in library complexity (Fig. S12).

Because our metrics are based on specific source annotation files, we also investigated the effect of alternative annotation source files. To illustrate, we recalculated exon:intron density ratios and pause indicies using UCSC RefSeq, Ensembl, and GENCODE gene set annotation files. While specific values per sample may have minor changes, as would be expected, the relationship between samples is consistent (Fig. S13).

## Conclusions

PEPPRO is an efficient, user-friendly PRO/GRO/ChRO-seq pipeline that produces novel, integral quality control plots and signal tracks that provide a comprehensive starting point for further downstream analysis. The included quality control metrics inform on library complexity, RNA integrity, nascent RNA purity, and run-on efficiency with theoretical and empirical recommended values (Fig. 9). PEPPRO is uniquely flexible, allowing pipeline users to serially align to multiple genomes, to select from multiple bioinformatic tools, and providing a convenient configurable interface so a user can adjust parameters for individual pipeline tasks. Furthermore, PEPPRO reads projects in PEP format, a standardized, well-described project definition format, providing an interface with Python and R APIs to simplify downstream analysis.

**Fig. 9:**
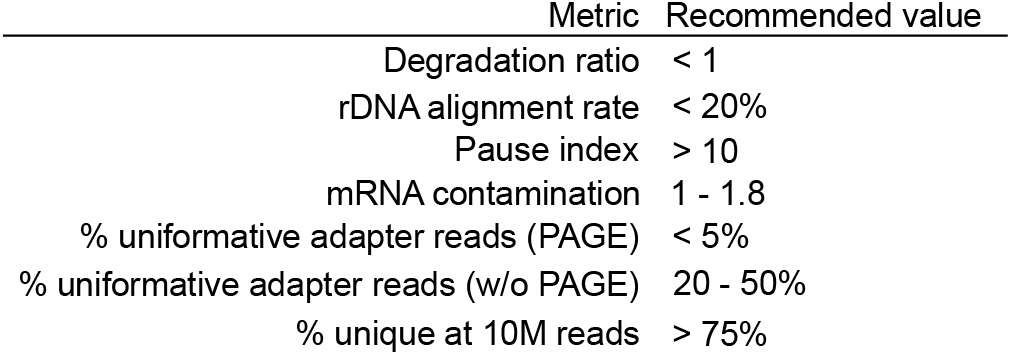
Recommendation table. Based on our experience processing both high- and low-quality nascent RNA libraries, these are our recommended values for high-quality PRO-seq libraries.

PEPPRO is easily deployable on any compute infrastructure, from a laptop to a compute cluster. It is thereby inherently expandable from single to multi-sample analyses with both group level and individual sample level quality control reporting. By design, PEPPRO enables simple restarts at any step in the process should the pipeline be interrupted. At multiple steps within the pipeline, different software options exist creating a swappable pipeline flow path with individual steps adaptable to future changes in the field. PEPPRO is a rapid, flexible, and portable PRO/GRO/ChRO-seq project analysis pipeline providing a standardized foundation for more advanced inquiries.

## Availability

Documentation on the Portable Encapsulated Project (PEP) standard may be found at pep.databio.org. `Refgenie` documentation and pre-built reference genomes are available at refgenie.databio.org. The PEP-PRO documentation, including links to an HTML report for the test samples, is hosted at peppro.databio.org, and source code is available at github.com/databio/peppro and archived under DOI 10.5281/zenodo.4542304.

## Methods

### Pipeline implementation

The PEPPRO pipeline is a python script (`peppro.py`) runnable from the command-line. PEPPRO provides restartability, file integrity protection, logging, monitoring, and other features. Individual pipeline settings can be configured using a pipeline configuration file (`peppro.yaml`), which enables a user to specify absolute or relative paths to installed software and parameterize alignment and filtering software tools. Required software includes several Python packages (cutadapt^25^, looper, numpy^26^, pandas^27^, pararead, pypiper, and refgenie^28^) and R packages (installed via the included PEPPROr R package) in addition to some common bioinformatics tools including bedtools^29^, bigWigCat^30^, bowtie2^31^, fastq-pair^32^, flash^33^, picard, preseq^20^, seqkit^18^, samtools^34^, seqtk, and wigToBigWig^30^. This configuration file will work out-of-the-box for research environments that include required software in the shell PATH, but may be configured to fit any computing environment and is adaptable to project-specific parameterization needs.

### Refgenie reference assembly resources

Several PEPPRO steps require generic reference genome assembly files, such as sequence indexes and annotation files. For example, alignment with `bowtie2` requires `bowtie2` indexes, and feature annotation to calculate fraction of reads in features requires a feature annotation. To simplify and standardize these assembly resources, PEPPRO uses *refgenie*. Refgenie is a reference genome assembly asset manager that streamlines downloading, building, and using data files related to reference genomes^28^. Refgenie includes recipes for building genome indexes and genome assets as well as downloads of pre-indexed genomes and assets for common assemblies. Refgenie enables easy generation of new standard reference genomes as needed. For a complete analysis, PEPPRO requires a number of refgenie managed assets. Those assets as defined by refgenie are: fasta, bowtie2 index, ensembl gtf, ensembl rb, refgene anno, and feat annotation. If building these assets manually, they separately require a genome fasta file, a gene set annotation file from RefGene, an Ensembl gene set annotation file in GTF format, and an Ensembl regulatory build annotation file. Finally, using PEPPRO with `seqOutBias` requires the additional refgenie tallymer index asset of the same read length as the data.

### Adapter-adapter ligation product abundance

Adapter-adapter ligation products show up in run-on libraries because there are two independent ligation steps. Sequencing these products is uninformative, and so there are several molecular approaches used to reduce their abundance in a sequencing library. All protocols include an inverted dT on the 3′ end of the 3′ adapter, and also do not phosphorylate the 5′ end of the 5′ adapter. Many protocols include a size-selection gel extraction step to purify the library from a prominent adapter-adapter ligation species.

PEPPRO calculates adapter-adapter ligation products directly from cutadapt output, and the default `-m` value for this step is the length of the UMI plus two nucleotides. Therefore, if RNA insertions fewer than three nucleotides in length are present in the library, these are treated as adapter-adapter ligation products.

### RNA insert size distribution and degradation

For both single and paired end data, the RNA insert size distribution is calculated prior to alignment. For single end data, the calculation is derived only from sequences that contain adapter sequence, which is output directly from `cutadapt`^25^. PEPPRO plots the inverse `cutadapt` report fragment lengths against the `cutadapt` fragment counts. If there is a known UMI, based on user input, that length is subtracted from reported `cutadapt` fragment lengths. As a consequence of this distribution, we can establish a measure of library integrity by evaluating the sum of fragments between 10-20 bases versus the sum of fragments between 30-40 bases in length. The higher this degradation ratio, the more degraded the library.

Paired end sequencing files often have shorter reads because a standard 75 base sequencing cartridge can be used for two paired end reads that are each 38 nucleotides in length. Therefore, many fewer of the reads derived from either end of the molecule extend into the adapter sequence. To address this issue, we incorporate a step that fuses overlapping reads using `flash`^33^. Therefore, if two paired end reads contain overlapping sequence, the reads are combined and the insert size is calculated directly from the fused reads and output directly from `flash`. This distribution is plotted identically to the single end reads and degradation is calculated in the same manner. This degradation ratio metric is uniform between single-end or paired-end libraries and is reported prior to any alignment steps, minimizing influences from extensive file processing or alignment eccentricities.

### Excluding size selection skews metrics

Recent PRO-seq protocols, including the H9 libraries we generated, exclude the PAGE size selection step that removes adapter-adapter ligation products^15^. Size selection can potentially bias against small RNA insertions. The previous two metrics: adapter-adapter abundance and degradation ratio are naturally skewed toward the undesirable range if libraries are constructed without size selection. Adapter abundance is skewed because the sole purpose of size selection is to remove the adapter species, but these uninformative reads are of minimal concern and can be overcome by increasing sequencing depth. Degradation ratio is skewed higher because the size selection is not perfect and insert sizes in the range of 10-20 are preferentially selected against relative to those in the 30-40 range. Therefore, while we provide recommendations for optimal degradation ratios, this metric is not necessarily comparable between library preparation protocols and a higher ratio is expected for protocols that exclude size selection.

### Removing UMI and reverse complementation

In a typical sequencing library, low library complexity is indicated by high levels of PCR duplicates. Conventional methods remove independent paired-end reads that map to the same genomic positions. This method works reasonably well for molecular genomics data sets with random nucleic acid cleavage. However, in PRO-seq, transcription start sites account for many of the 5′ RNA ends and polymerases pause downstream in a focused region^4^. Consequently, independent insertions with the same end points are common, especially in the promoter-proximal region. To solve this, PRO-seq protocols incorporate a unique molecular identifier (UMI) into the 3′ adapter to distinguish between PCR duplicates and independent insertions with shared ends. PEPPRO removes PCR duplicates only if UMIs are provided.

Following the removal of PCR duplicates, the UMI is trimmed. For run-on experiments where the sequencing primer sequences the 3′ end of the original RNA molecule, reverse complementation is performed. As only the first read contains a UMI in paired-end experiments, the second reads skip UMI trimming. Both steps are performed using either `seqtk` (https://github.com/lh3/seqtk) or fastx (https://github.com/agordon/fastx_toolkit), depending on user preference. Because reads are processed uniquely for first and second reads in a paired-end experiment, reads must be re-paired prior to alignment. PEPPRO uses the optimized implementation `fastq-pair`^32^ to re-pair desynchronized read files.

### Serial alignments

Following re-pairing, or starting from processed single-end reads, PEPPRO performs a series of preliminary, serial alignments (prealignments) before aligning to the primary reference using `bowtie2`^31^. As a significant portion of nascent transcription includes rDNA, PEPPRO defaults to initially aligning all reads to the human rDNA sequence. Not only does this remove rDNA reads from downstream analysis, it improves computational efficiency by aligning the largest read pool to a small genome and reduces that read pool for subsequent steps. The user can specify any number of additional genomes to align to prior to primary alignment, which may be used for species contamination, dual-species experiments, repeat model alignments, decoy contamination, or spike-in controls. For serial alignments, `bowtie2` is run with the following parameters `-k 1-D 20 -R 3 -N 1 -L 20 -i S,1,0.50`, where we are interested primarily in quickly identifying and removing any reads that have a valid alignment to the serial alignment genome (-k 1 parameter). These settings are easily adjusted in the pipeline configuration file (`peppro.yaml`).

Subsequent to these serial alignments, remaining reads are aligned to the primary genome. Primary genome alignment uses the `bowtie2 --very-sensitive` option by default and sets the maximum paired-end fragment length to 2000. The goal with primary alignment is to identify the best valid alignment for reads, sacrificing speed for accuracy. Following primary alignment, low-quality reads are removed using `samtools view –q 10`. As with the initial prealignments, these parameters can be customized by the user in the pipeline configuration file (`peppro.yaml`). Alignment statistics (number of aligned reads and alignment rate) for all serial alignments and primary alignments are reported. For the primary alignment, PEPPRO also reports the number of mapped reads, the number removed for quality control, the total efficiency of alignment (aligned reads out of total raw reads), and the read depth. Prior to further downstream analysis, paired-end reads are split into separate read alignment files and only the first read is retained for downstream processing. For both paired-end and single-end experiments, this aligned read file is split by strand with both plus and minus strand aligned files further processed.

### Processed signal tracks

Following read processing, alignment, strand separation, and quality control reporting, aligned reads are efficiently converted into strand-specific bigWig files by default. For PRO-seq and similar protocols, reads are reported from the 3′ end and may optionally be scaled by total reads. PEPPRO may alternatively use `seqOutBias`^19^ to correct enzymatic sequence bias. Bias is corrected by taking the ratio of genome-wide observed read counts to the expected sequence based counts for each k-mer^19^. K-mer counts take into account mappability at a given read length using Genome Tools’ `Tallymer` program^35^. Correcting for enzymatic bias can be important as bias from T4 RNA Ligase used in PRO-seq protocols can yield erroneous conclusions^19^. As such, we recommend using `seqOutBias` for bias correction when analyzing a typical PRO-seq library. Bias correction is especially important when plotting composite profiles over sequence features. Strand specific bigWigs may be visually analyzed using genomic visualization tools and provide a unified starting point for downstream analyses. For example, output bigWig files can be directly loaded into `dREG` to identify regulatory elements defined by bidirectional transcription^1^.

### Exon-intron ratio plots

PEPPRO provides an mRNA contamination histogram for quick visual quality control, and a BED format file containing gene by gene exon:intron ratios for detailed analysis. To calculate this metric, PEPPRO utilizes annotation files derived from UCSC RefSeq gene files. Because promoter-proximal pausing inflates these ratios, PEPPRO excludes the first exon from the calculation. Otherwise, the reads per kilobase per million mapped reads (RPKM) is calculated for all exonic and intronic sequences on a gene by gene basis. Then, the ratio of exon RPKM to intron RPKM is determined *for every gene*. The overall measure, the mRNA contamination metric, is the median of all genic exon:intron density ratios.

### Pause index

Pause indices are calculated as the ratio of read density in the promoter proximal region versus read density in the gene body. To calculate these values, PEPPRO utilizes annotation files derived from Ensembl gene set files. Pause indices can vary widely depending on the defined pause window and how a pause window is determined (i.e. relative to a TSS or the most dense window proximal to a TSS). PEPPRO defines the density within the pause region as the single, most dense window +20-120 bp taken from all annotated TSS isoforms per gene. This is necessary as some genes contain multiple exon 1 annotations and because this region is where most polymerase pausing occurs, PEPPRO identifies the predominant exon 1, based on density, and calculates the pause index using this window density. This means that for genes with multiple TSSs, we define the pause window as the region +20-120 bases from *each* identified TSS per gene. We determine the read density at every annotated pause window per gene, and identify the predominant, singular pause window as the pause window with the greatest density. This singular pause window is used to calculate the pause index for that gene. The corresponding gene body is defined as the region beginning 500 bp downstream from the predominant TSS to the gene end. We found that lowly expressed genes represent a significant portion of genes with a low pause index. At low sequencing depth, these lowly expressed genes experience greater dropout and fluctuation in pause index calculation, skewing the metric upwards at low depth. To address this, we restrict the pause index calculation to the upper 50th percentile of genes by expression, which eliminates the variability due to depth. Finally, PEPPRO plots the distribution of pause indices for each remaining gene in a histogram and provides a BED-formatted file containing each gene’s pause index for more detailed analyses.

### PRO-seq experiments

H9 PRO-seq experiments were conducted as described previously^15^. The HDACi-treated samples were incubated with 200nM romidepsin for 60 minutes prior to harvesting. The control “untreated” samples were treated with DMSO for 60 minutes. We have included these samples as a test to demonstrate differential expression analysis using PEPPRO. They also provide additional example libraries for the metrics in general, and unexpectedly, show significant differences in pause index upon treatment.

### Synthetic experiments

Synthetic sequencing depth variant libraries were constructed for single-end and paired-end PRO-seq libraries using either the K562 PRO-seq (GSM1480327) or H9 PRO-seq 2 (GSM4214081) as source libraries, respectively. For K562 PRO-seq subsamples, `seqtk sample -s99` was called on the raw fastq files to generate libraries between 2-10%, in 2 percent increments, and between 10-100%, in 10 percent increments. For the H9 PRO-seq libraries, `seqtk sample -s99` was called on the raw fastq files to produce libraries between 10-100%, in 10 percent increments. Lower percentage K562 PRO-seq libraries were generated to yield libraries of total size comparable to low percentage H9 PRO-seq libraries.

RNA-seq spike-in libraries were also produced using the command `seqtk sample -s99` on raw fastq files using combinations of the K562 PRO-seq library utilized prior and a corresponding K562 RNA-seq library (GSM765405). RNA-seq libraries were sampled between 10-100%, in 10 percent increments, and concatenated with the sampled K562 PRO-seq libraries to generate mixed libraries composed of 0-100% RNA-seq.

Low complexity libraries were similarly constructed. Thirty million total read libraries were generated by using `seqtk sample -s99` on the H9 PRO-seq 2 library and sampling at 50, 80, 90, 92, 94, 96, 98, and 100%. At each percentage of original H9 PRO-seq 2 library sample, the remainder represents duplicates of the original raw reads composing the opposite percentage, producing libraries with varying levels of duplicated reads.

## Funding

This work was supported by the National Institute of General Medical Sciences grants GM128635 (MJG) and GM128636 (NCS). JPS was supported by the institutional training grant GM008136. ABD was supported by institutional training grants LM012416 and GM007267.

## Declarations

All authors declare no competing interests.

## Supplemental text

### Gene counts table

`PEPPRO` provides a project level counts table that simplifies downstream analyses. Here, we import the PEPPRO

~~~
project counts table and construct a DESeq data set in a few lines of code.
# 1. Load the PEPPRO R package.
library(PEPPROr)
# 2. Load the PEP project using the project configuration file.
prj = Project(“peppro_paper.yaml”)
# 3. Load the project gene counts table.
counts = read.csv(file.path(paste0(config(prj)$metadata$output_dir,
              “/summary/PEPPRO_countData.csv”)))
# 4. Only keep the H9 untreated or H9 HDAC inhibitor treated samples.
counts = counts[,c(“geneName”, “H9_PRO-seq_1”, “H9_PRO-seq_2”, “H9_PRO-seq_3”,
          “H9_treated_PRO-seq_1”, “H9_treated_PRO-seq_2”,
          “H9_treated_PRO-seq_3”)]
# 5. Convert the counts table to a matrix by removing the gene name column.
count_matrix = as.matrix(counts[,-”geneName”])
# 6. Set the rownames of the matrix object to be the gene names.
rownames(count_matrix) = counts$geneName
# 7. Create a data.frame that defines the sample information.
coldata = data.frame(condition=c(rep(“untreated”, 3), rep(“treated”, 3)))
# 8. Set the rownames of the sample information data.frame to match the counts matrix. rownames(coldata) = colnames(count_matrix)
# 9. Load the DESeq2 package.
library(“DESeq2”)
# 10. Create a DESeq data set from our counts matrix and the sample information data.frame.
dds = DESeqDataSetFromMatrix(countData = count_matrix,
               colData = coldata,
               design = ∼ condition)
~~~

## Supplemental figures

**Fig. S1:**
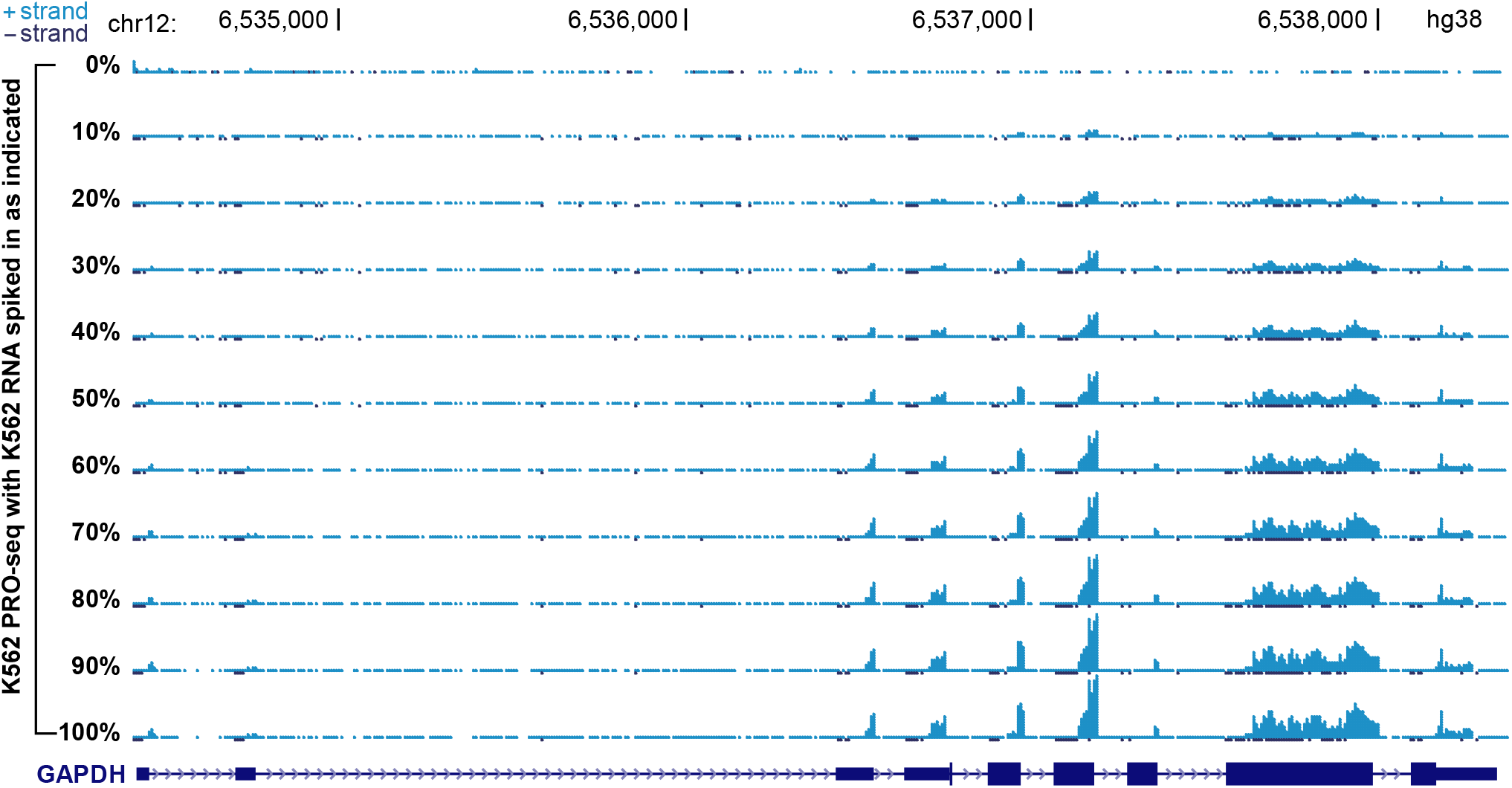
K562 RNA-seq spike-in signal tracks show increasing exonic coverage. GAPDH exonic coverage is enriched as the percentage of RNA-seq reads increases, and is visualized particularly well at exons 6 and 8. Each sample library is composed of 70M total reads.

**Fig. S2:**
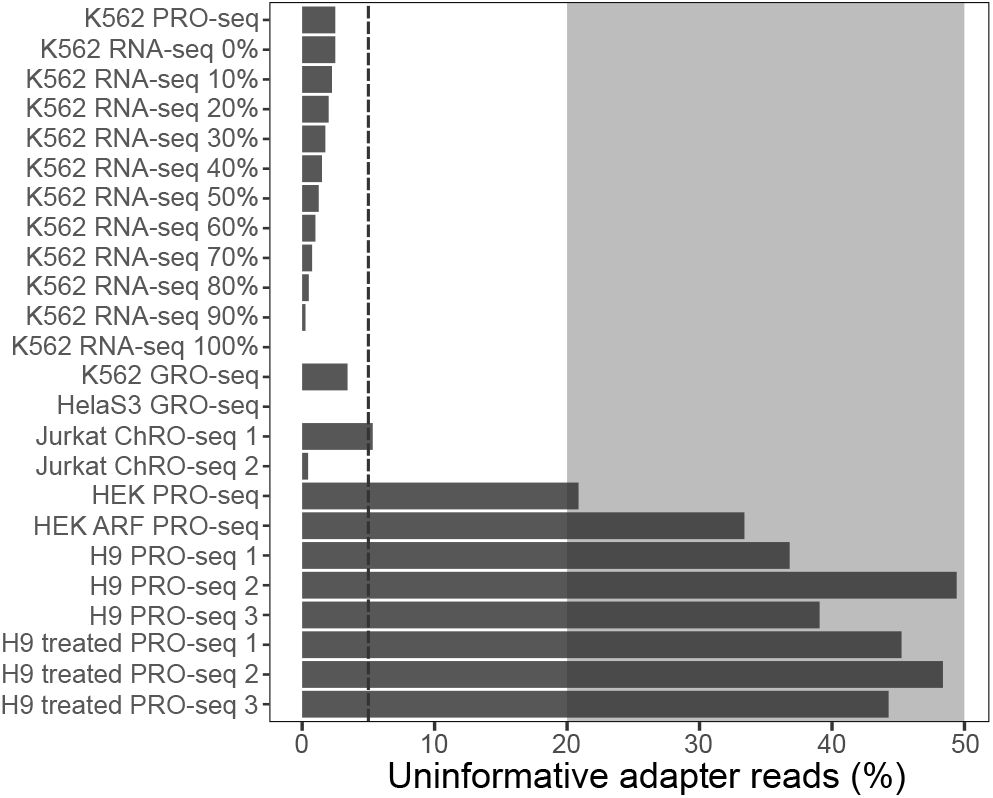
Percentage of uninformative adapter reads following adapter removal for test set samples. The HEK and H9 libraries contain more adapter-adapter reads because PAGE-mediated size selection was excluded from the protocol (Values below the dashed line are generally recommended for PAGE-purified libraries. Shaded region represents the recommended abundance of adapter reads for libraries without PAGE purification.)

**Fig. S3:**
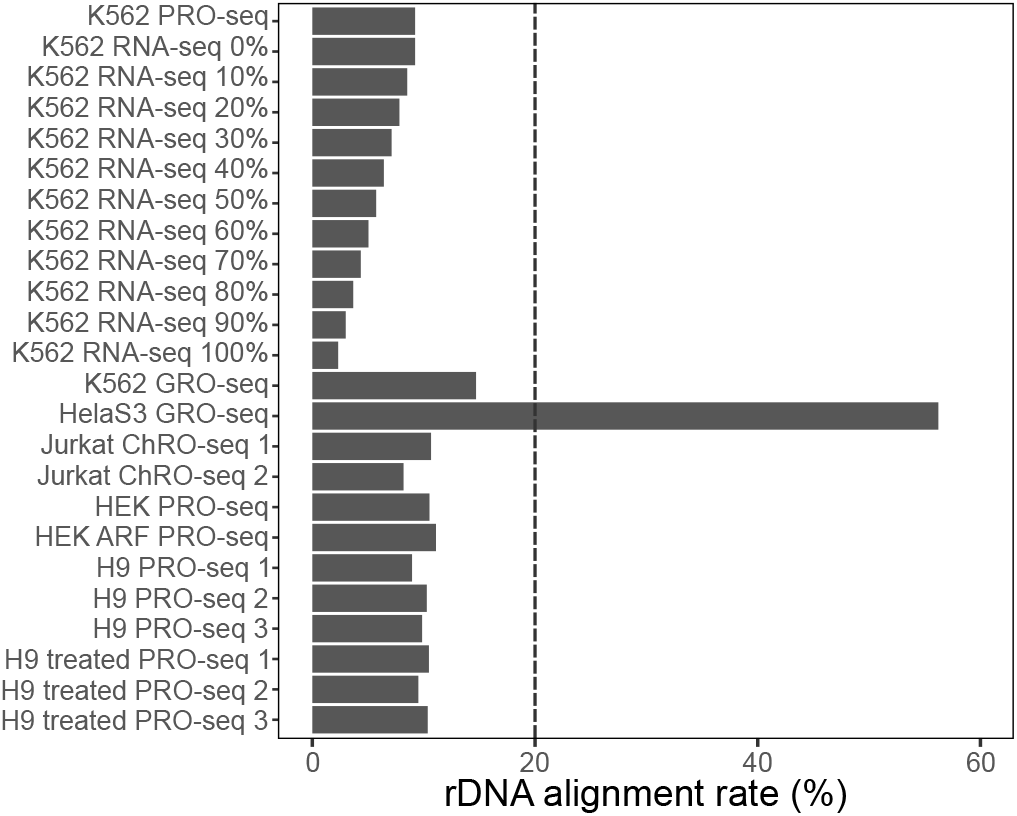
Ribosomal DNA alignment rates for test set samples. The HelaS3 GRO-seq sample is highly enriched for ribosomal RNA transcripts compared to other test samples. Values below the dashed line are recommended.

**Fig. S4:**
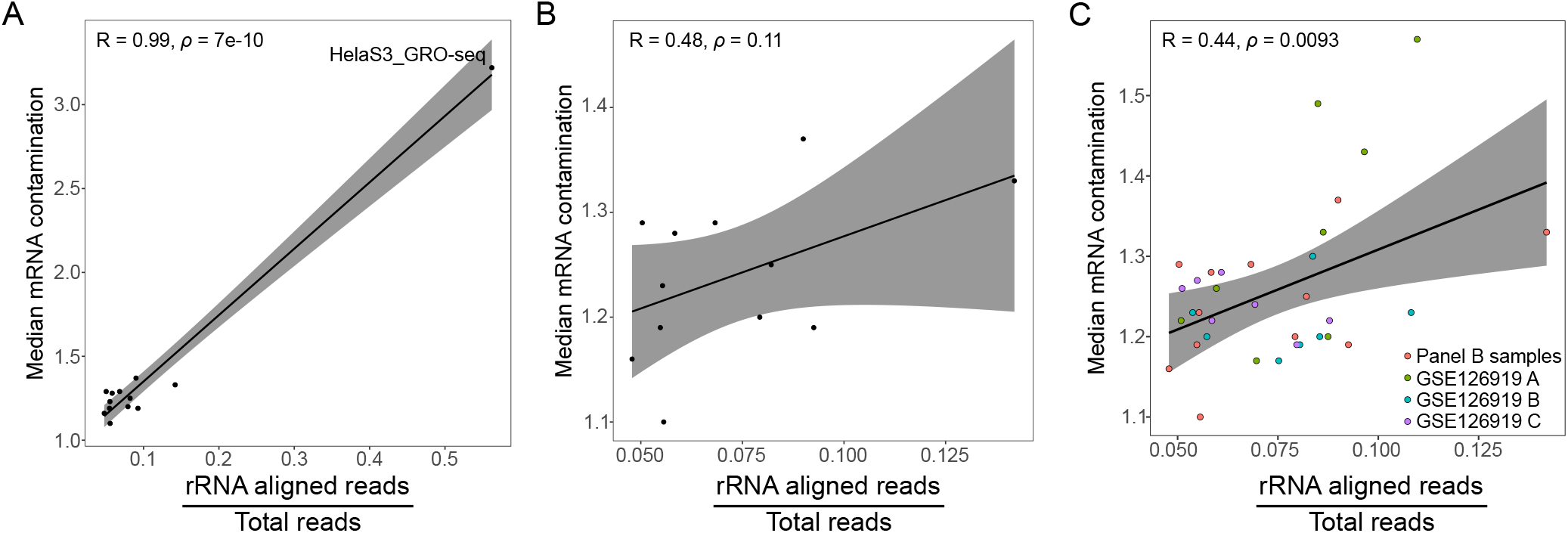
Abundance of rDNA to total reads is correlated with mature RNA contamination. Correlation plot between the measure of mRNA contamination (median exon:intron density) and the ratio of rDNA aligned reads to total reads for: A) all primary samples (*excludes RNA-seq spike-in experiment due to ribosomal depletion inherent in RNA-seq library preparation), B) primary samples excluding the known outlier HelaS3 GRO-seq sample, C) all samples in panel B and all non-redundant samples from GSE126919 including three cellular subclones (A, B, and C) to demonstrate possible differences due to cell lines. Test for association determined with Pearson’s product moment correlation coefficient.

**Fig. S5:**
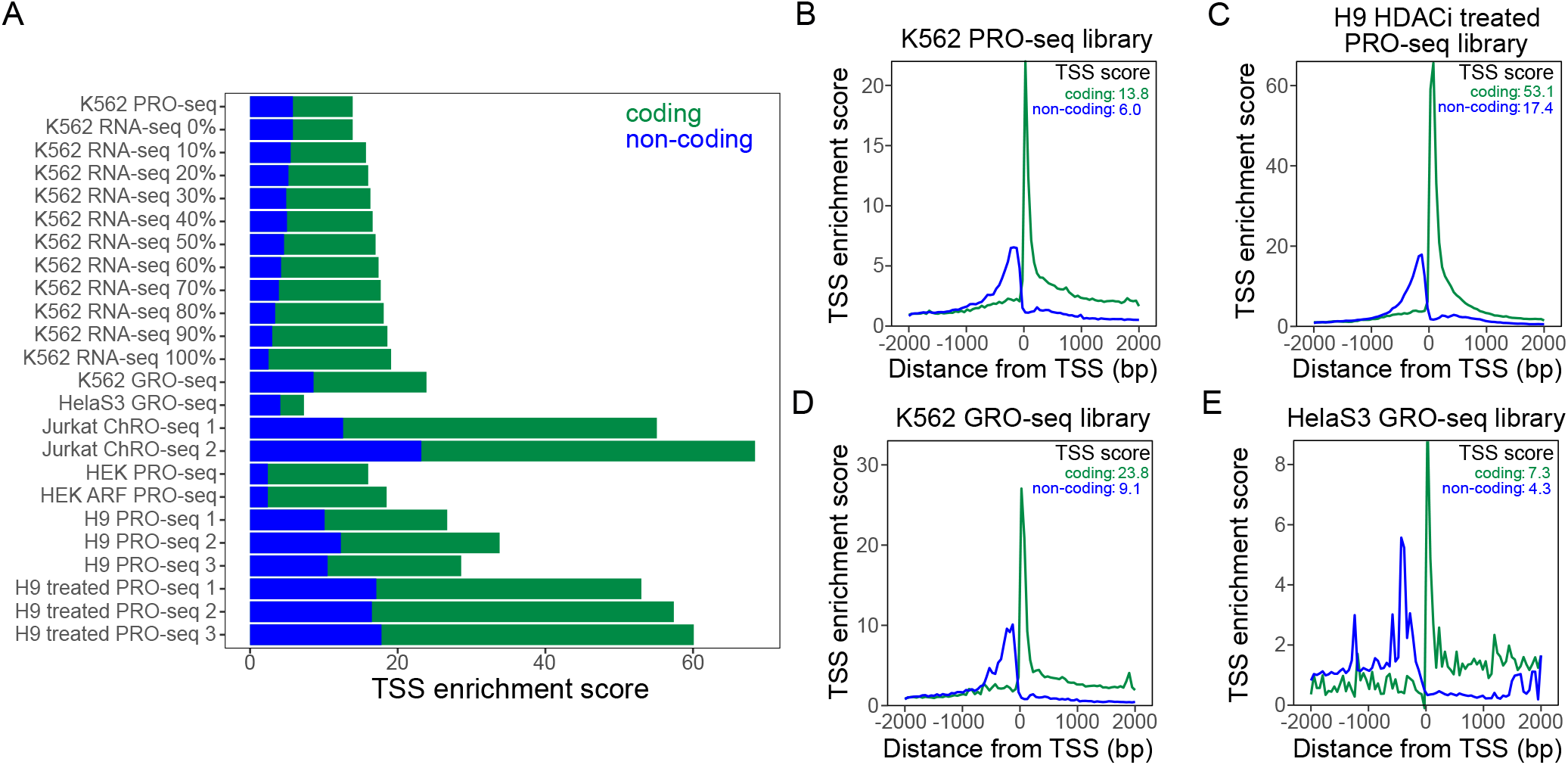
TSS enrichment. A) TSS enrichment scores for test set samples. B) Representative high-quality PRO-seq TSS enrichment plot. C) TSS enrichment plot in romidepsin treated PRO-seq library. D) Representative high quality GRO-seq TSS enrichment plot E) Representative example of lower quality GRO-seq TSS enrichment plot.

**Fig. S6:**
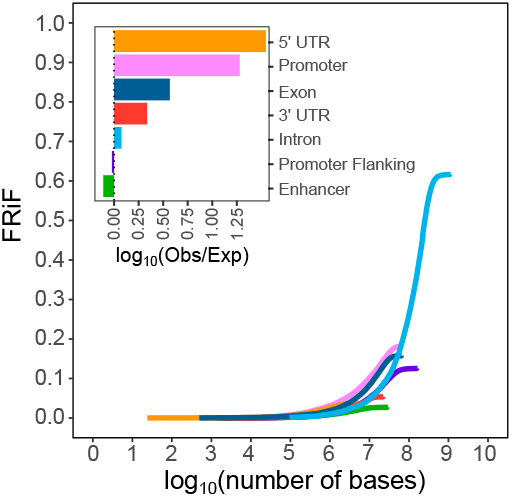
Fraction of Reads in Features in ChRO-seq. Cumulative FRiF and FRiF (inset) plots for example Jurkat ChRO-seq 1 library test sample shows increased enrichment of promoter sequences.

**Fig. S7:**
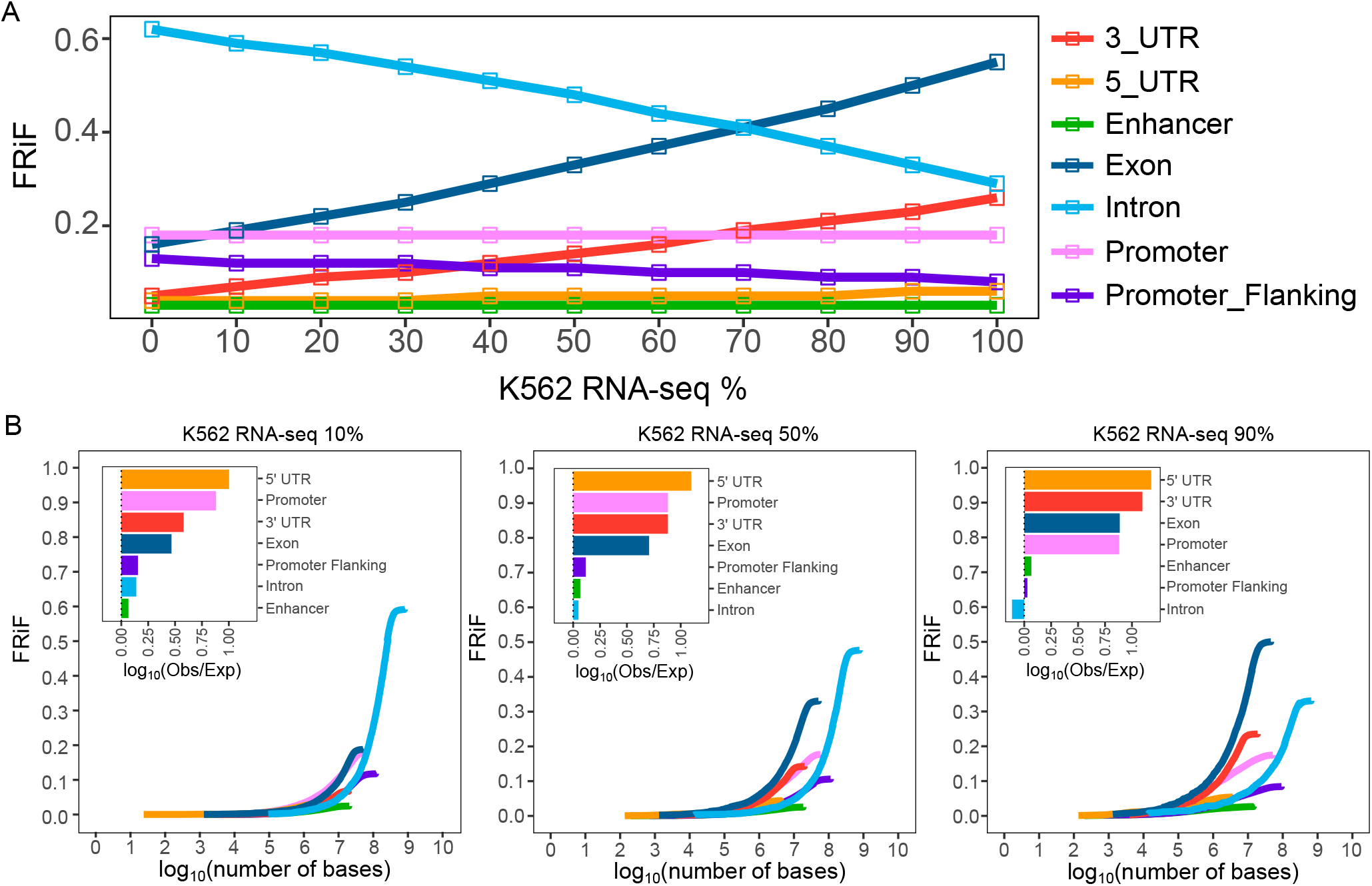
RNA-seq spike-in shows how FRiF changes with mRNA contamination. A) Increasing percentages of RNA-seq spike-in lead to changes in the fraction of reads in features (FRiF). B) Cumulative FRiF plots at 10%, 50%, and 90% RNA-seq spike-in. Plot insets represent the expected versus observed fraction of reads in genomic features. Each spike-in library contains 70 million total reads.

**Fig. S8:**
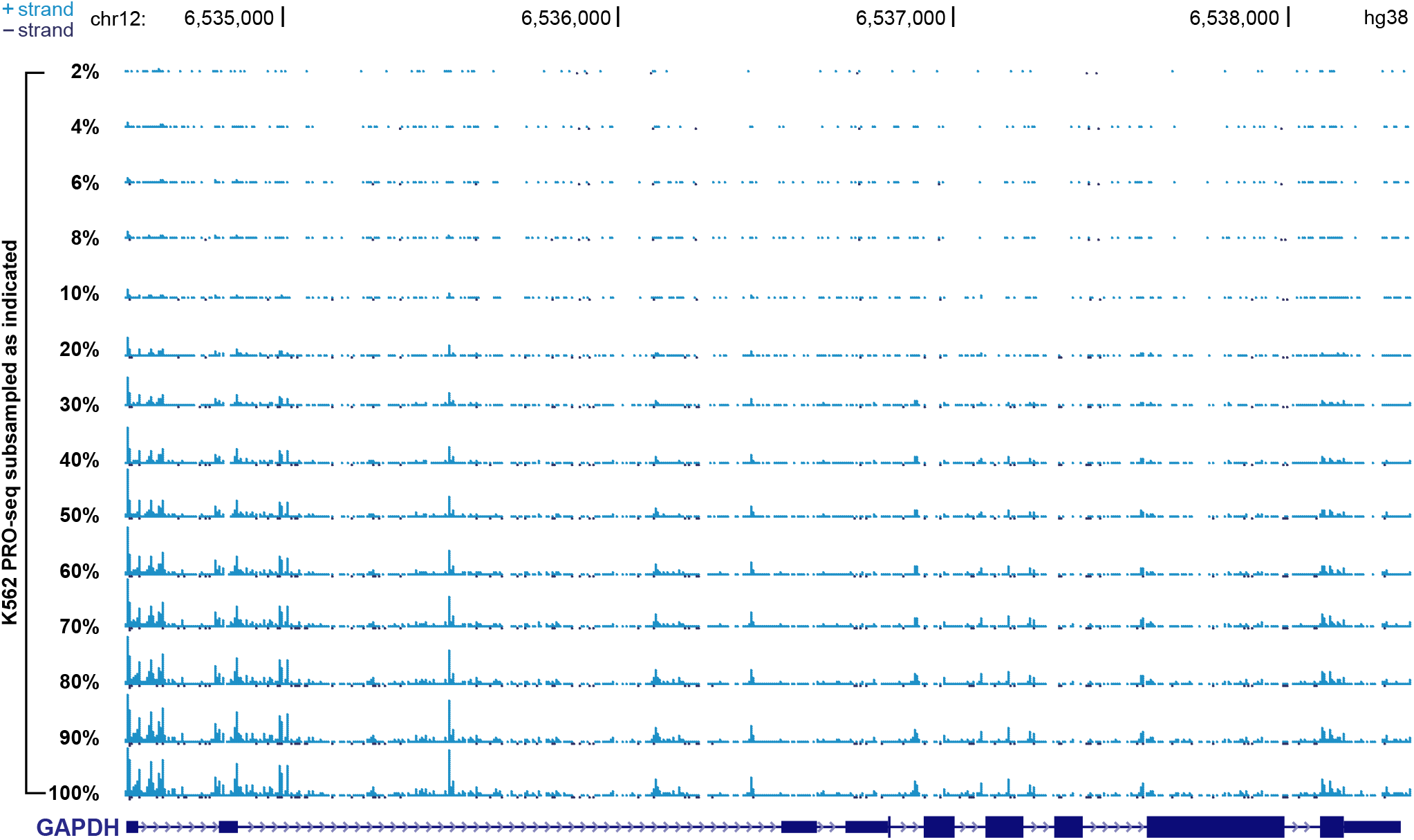
K562 PRO-seq signal tracks show increasing coverage with depth. Incrementally subsampled K562 PRO-seq library signal tracks display reduced relative coverage at a representative locus (GAPDH).

**Fig. S9:**
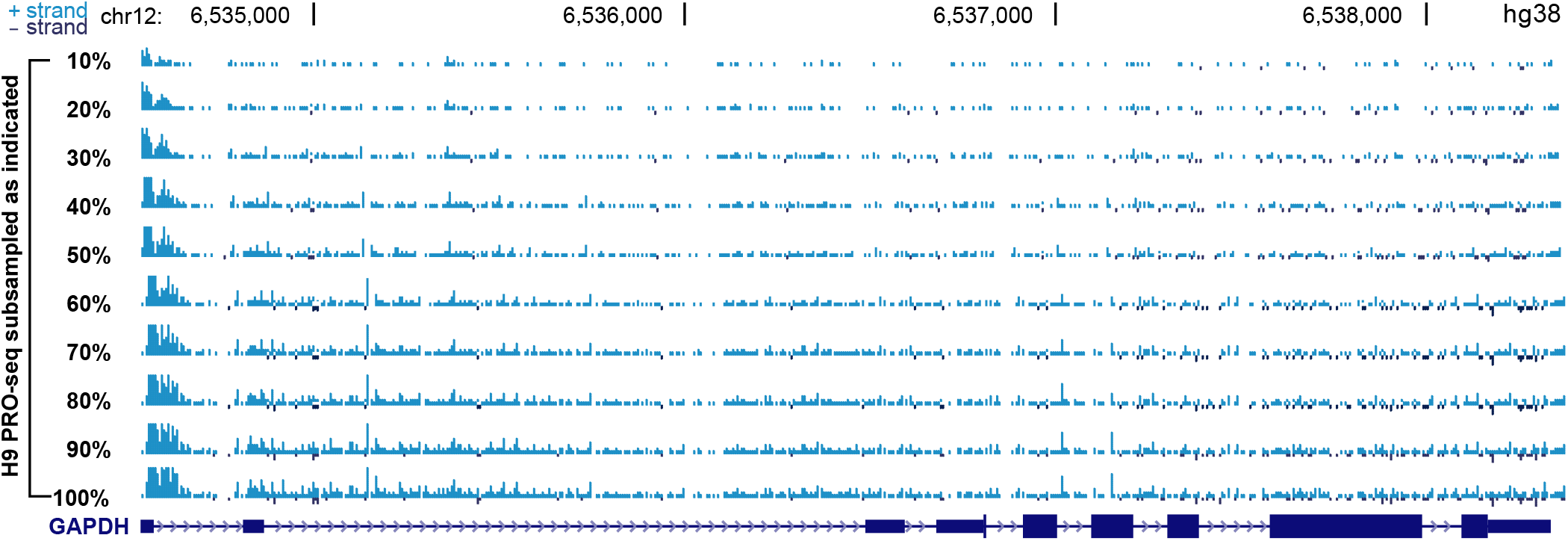
H9 PRO-seq 2 signal tracks show increasing coverage with depth. Incrementally subsampled H9 PRO-seq 2 library signal tracks display reduced relative coverage at a representative locus (GAPDH).

**Fig. S10:**
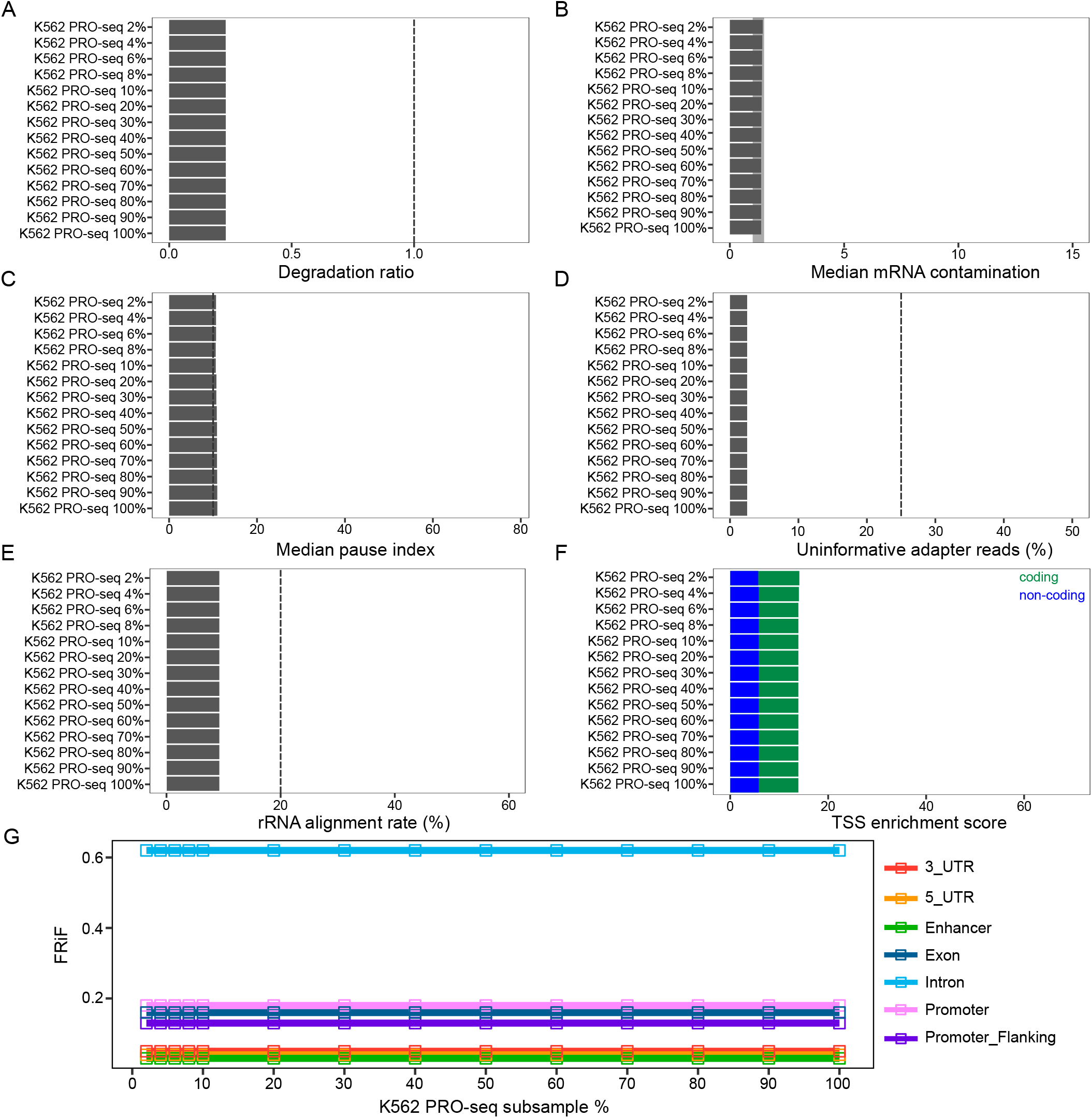
QC metrics are not affected by sequencing depth in subsampled K562 PRO-seq. Using subsampled K562 PRO-seq data, we show how various metrics behave across a spectrum of sequencing depths: A) Degradation ratio, B) mRNA contamination, C) Pause index, D) the percentage of uninformative adapter reads, E) the rDNA alignment rate, F) and the TSS enrichment scores are unaffected by sequencing depth. G) The FRiF and cumulative FRiF is unaffected by sequencing depth. The complete K562 PRO-seq library (100%) contains approximately 497 million reads.

**Fig. S11:**
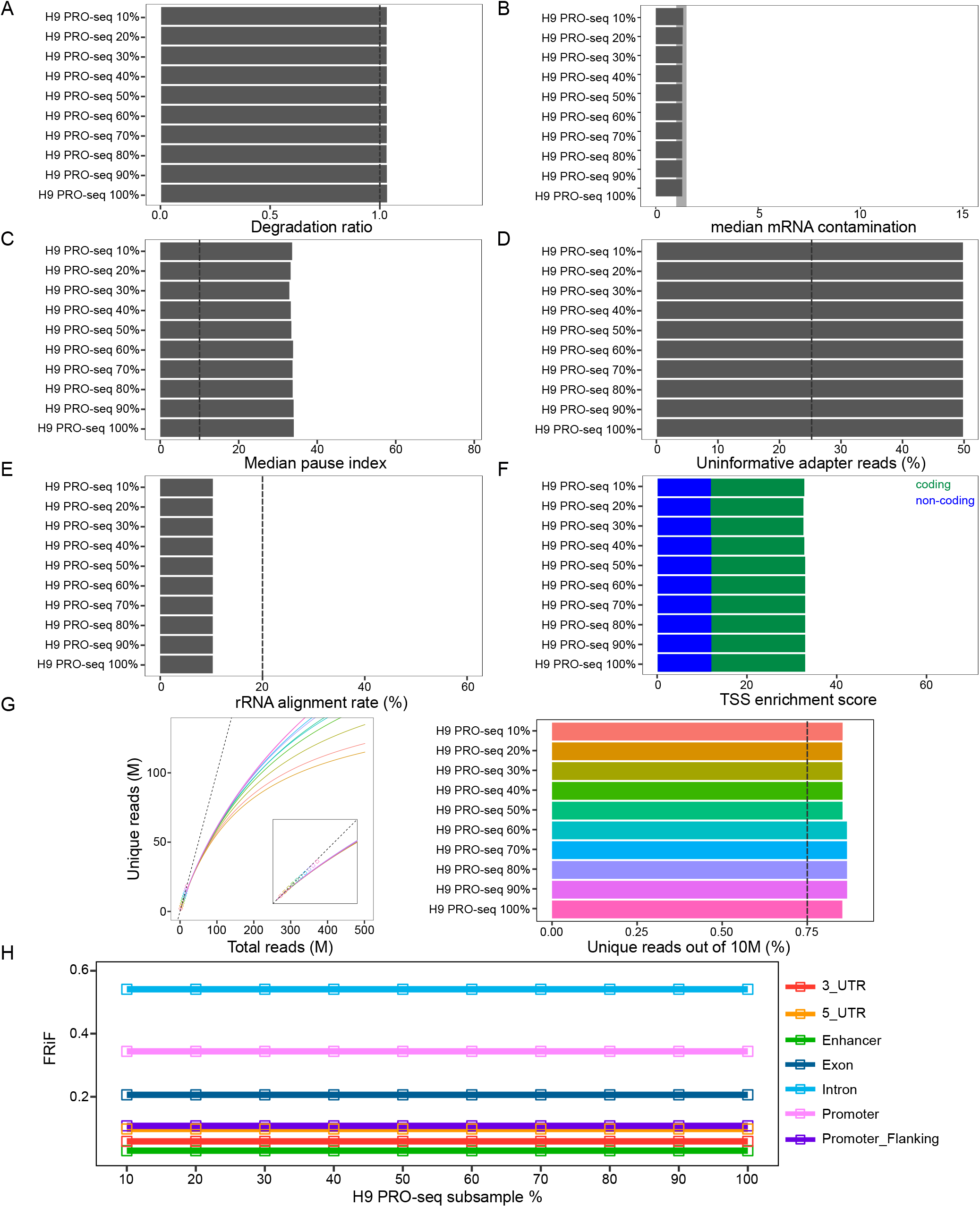
QC metrics are not affected by sequencing depth in subsampled H9 PRO-seq. Using subsampled H9 PRO-seq data, we show how various metrics behave across a spectrum of sequencing depths: A) Degradation ratio, B) mRNA contamination, C) Pause index, D) the percentage of uninformative adapter reads, E) the rDNA alignment rate, F) and the TSS enrichment scores are unaffected by sequencing depth. G) Library complexity traces plot the read count versus externally calculated deduplicated read counts. Deduplication is a prerequisite, so these plots may only be produced for samples with UMIs. Inset zooms to region from 0 to double the maximum number of unique reads. The position of curves in the left panel at a sequencing depth of 10 million reads (dashed line represents minimum recommended percentage of unique reads). H) The FRiF and cumulative FRiF is unaffected by sequencing depth. The complete H9 PRO-seq library (100%) contains approximately 116 million reads.

**Fig. S12:**
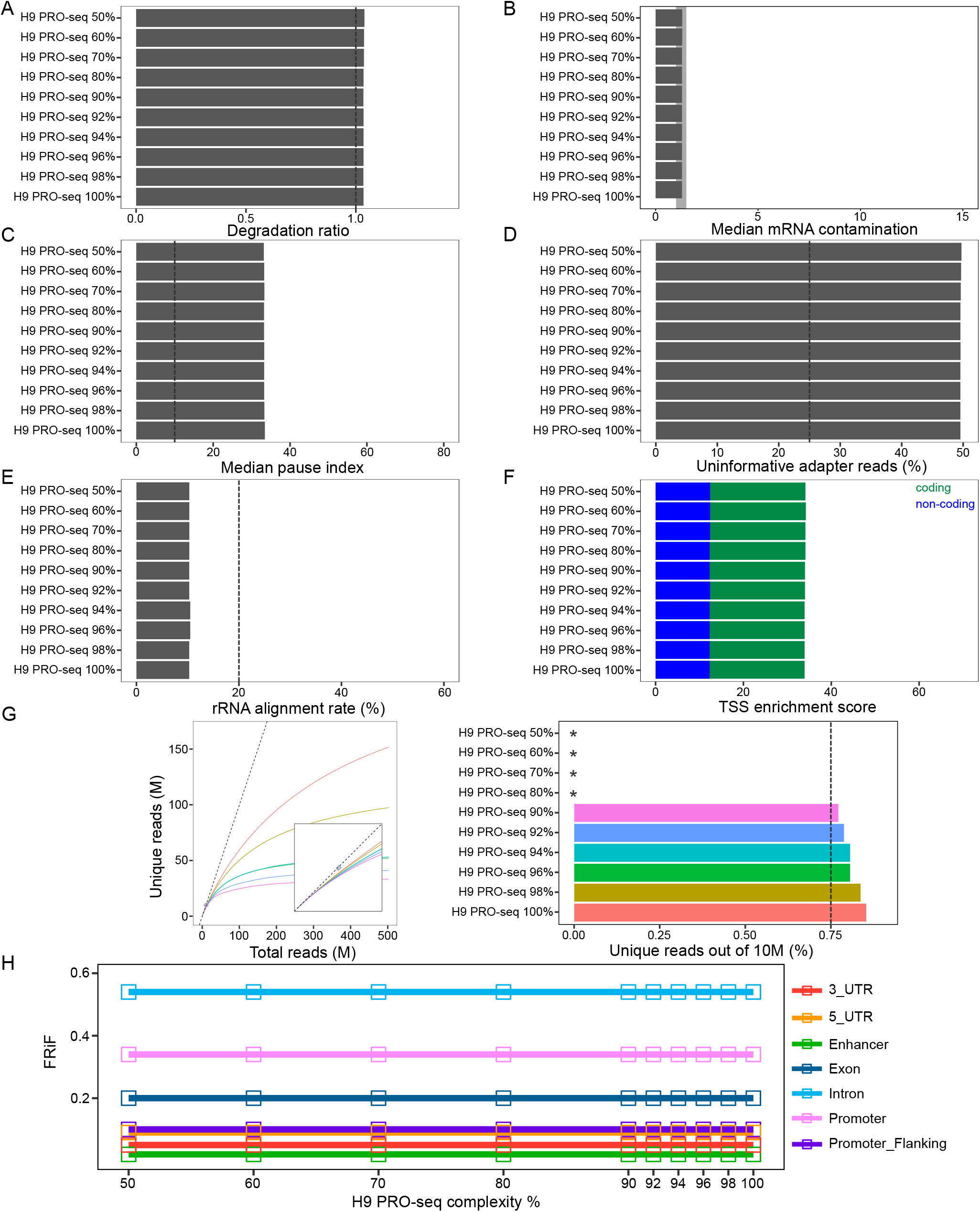
QC metrics are not affected by low library complexity. Using a synthetic set of libraries, we show how various metrics behave across a spectrum of complexity: A) Degradation ratio, B) mRNA contamination, C) Pause index, D) the percentage of uninformative adapter reads, E) the rDNA alignment rate, F) and the TSS enrichment scores are unaffected by low complexity. G) Library complexity traces plot the read count versus externally calculated deduplicated read counts. Deduplication is a prerequisite, so these plots may only be produced for samples with UMIs. Inset zooms to region from 0 to double the maximum number of unique reads. The right panel represents the position of curves in the left panel at a sequencing depth of 10 million reads (dashed line represents minimum recommended percentage of unique reads). *Libraries with less than 90% uniqueness could not be extrapolated due to saturation. H) The FRiF and cumulative FRiF is unaffected by low complexity. Each library contains 30 million total reads.

**Fig. S13:**
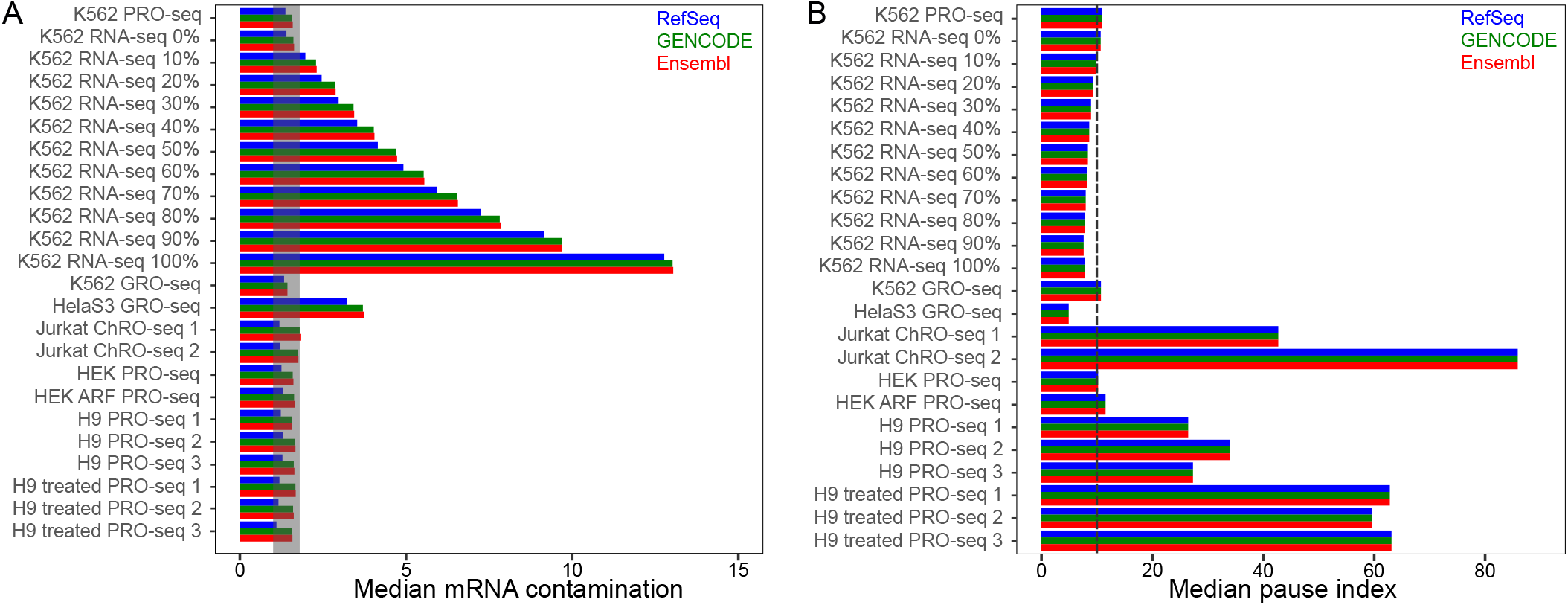
Alternate annotation sources do not affect mRNA contamination and pause index. A) The mRNA contamination metric and B) the pause index metric are robust across annotations.

## Notes

http://peppro.databio.org

https://github.com/databio/peppro

### Competing Interest Statement

The authors have declared no competing interest.

### Summary of Updates

New supplementary figure, text clarifications.

